# Metagenomic Discovery of Neutral Lipid Metabolism Pathways in the Arctic Ocean Microbiomes Suggests a Potential New Role in Survival and Oceanic Carbon Cycling

**DOI:** 10.64898/2026.07.24.740503

**Authors:** Thomas Grevesse, Susan McLatchie, Vera E. Onana, David A. Walsh

## Abstract

The Arctic Ocean microbiomes experience extreme seasonal fluctuations in light, nutrient availability, and organic carbon supply. In this environment, neutral lipid storage may provide a key survival strategy. Here, we investigated the diversity, distribution, and ecological role of neutral lipid metabolism in Arctic microbiomes using metagenome-resolved analyses and global ocean comparisons. Arctic photic-zone microbiomes were strongly enriched in triacylglycerol (TAG) biosynthesis genes relative to other oceans, primarily due to picoeukaryotic phytoplankton, including the ecologically dominant *Micromonas* and *Bathycoccus*. In contrast, prokaryotic communities exhibited diverse TAG-degrading taxa and fatty acid transport systems, supporting a previously unrecognized lipotrophic bacterial guild exploiting phytoplankton-derived lipids as carbon and energy sources. Genome-resolved analyses further revealed distinct bacterial lipid-storage strategies: TAG-producing taxa preferentially encoded fatty acid uptake and carbohydrate utilization pathways, whereas polyhydroxyalkanoate-producing taxa were associated with aromatic compound degradation, linking terrestrial organic matter to lipid storage. Our results expand the known diversity of marine microbes capable of neutral lipid metabolism and identify microbial lipid cycling as a previously overlooked component of the Arctic carbon cycle. We propose that neutral lipid storage and turnover support microbial survival through the polar night while enhancing carbon transfer within Arctic microbial food webs under ongoing climate change.

## Introduction

The microbial communities of the Arctic Ocean are faced with intense seasonal changes in energy, carbon and nutrient availability. During spring, the nutrients released by ice melt and river discharge, combined with longer light days, create a simultaneous abundance of CO_2_, energy and nutrients for microbial phytoplankton, triggering blooms at the surface mixed layer^1^. In summer and fall, phytoplankton lack nutrients that have been depleted during spring blooms but CO_2_ and light energy are still abundant and available with the long days and absence of ice. In winter, the Arctic Ocean receives very little light due to short days and a thick ice-layer, limiting primary production. The Arctic Ocean heterotrophs also face intense seasonal variations in carbon, nutrient and energy. The spring phytoplankton blooms, combined with riverine discharge, constitute a pulse of organic matter for Arctic Ocean heterotrophs. The prokaryotic heterotrophs must then live on diminishing primary produced organic matter (OM) as summer and fall progress^2^ and finally on the recycling of the leftover stock of primary-produced OM and decaying living matter during the polar night^3^.

In a context of strong variations in energy and nutrients, it may be an advantageous life strategy for Arctic Ocean microbial communities to accumulate chemical resources in periods of abundance in prevision of periods of scarcity, a process called storage^4^. Among the classes of compounds used to store organic carbon (C), lipids have the highest energy density (up to 9.3 kcal/g) compared to proteins and carbohydrates (4.1 kcal/g)^5^. Lipid storage is therefore a common trait across the domains of life^6–8^. Neutral lipids (NLs) are the common storage form of lipids, and can be differentiated from the charged lipids that comprise cellular membranes^9^. There are three NLs produced by eukaryotes: triacylglycerols (TAGs), steryl esters (SEs), and wax esters (WEs). All are synthesized through the esterification of a fatty acid (FA) with a second moiety: diacylglycerol is used to form TAGs, a sterol to form SEs, or a fatty alcohol to form WEs^7^. Although TAG biosynthesis, and to a lesser extent WE and SE^8,10,11^ biosynthesis, has been observed in bacteria, polyhdroxyalkanaoates (PHAs) are a more widespread bacterial energy storage compound^8^. The shortest PHA, polyhydroxybutyrate (PHB) is produced from the condensation of two acetyl-CoA molecules into a 3-hydroxy-butyryl-CoA (3HB) followed by its polymerization^12^. Alternatively short molecules such as glutamic acid, butyric acid or succinic acid can be transformed into 3HB and polymerized to PHB^13^. Longer chain PHAs are polymerized following two routes: (i) from 3-hydroxyacyl-CoA intermediates diverted from β-oxidation when FAs are used as carbon sources^13,14^ and (ii) from 3-hydroxyacyl-CoA obtained through *de novo* FA biosynthesis in the presence of glucose, acetate or ethanol as carbon sources^13^.

The metabolism of NLs plays important ecological roles for marine phytoplankton. NLs, especially TAGs are stored during the day and consumed during the night, representing diel oscillations amounting to almost a quarter of the total daily primary productivity^15^. NLs, in the form of lipid droplets in the cell also act as lipid reservoir to alleviate the lipotoxicity of lipid intermediates (free FAs, DAGs, etc) but also as a pool that delivers stored NL for energy production, membrane homeostasis and growth^16^. NLs are also involved in external stressor response and alleviation by recycling membrane lipids^17^. NL production, and specifically TAG, has been widely reported in numerous ocean phytoplankton species^15,18^. In polar environments, the use of accumulated NLs, especially TAGs, coupled with reduced metabolism has been demonstrated as a strategy for some phytoplankton species to overwinter during the dark polar night both in the Arctic^19^ and Antarctic^20^ Oceans. NL production and accumulation are generally favored when C and energy sources are abundant and other nutrients (N, P, etc.) are limited^21,22^. The conditions of the Arctic summer and fall, with long days providing ample energy from light and waters depleted of nutrients by the spring blooms^23^ are therefore ideal for the accumulation of NLs in phytoplankton. In addition, the increased stratification of the Arctic Ocean, preventing nutrients resupply to the photic zone^23^ may further favor NL accumulation in phytoplankton species. In the Arctic Ocean however, only a few studies explicitly identified the production of NLs in phytoplankton isolates ^19^. And despite the important ecological role that NL metabolism may play for the Arctic phytoplankton, we still don’t know the phylogenetic distribution of NL metabolism across Arctic phytoplankton.

The production and degradation of NLs as a survival strategy for the Arctic Ocean phytoplankton may contribute importantly to the food web and C cycle of the Arctic Ocean. The Arctic Ocean phytoplankton communities are dominated by eukaryotic algae, with a general absence of Cyanobacteria^24^. In addition, picoeukaryotes are active and often dominate phytoplankton populations of the Arctic, especially in oligotrophic regions such as the Canada Basin^25,26^ and thrive under changing conditions associated with climate change^27^. Under nutrient limitation, eukaryotic phytoplankton store energy in the form of neutral lipids and carbohydrates^28^, while Cyanobacteria favour carbohydrates. Nutrient limitation of oligotrophic waters also favors smaller picoeukaryotes over larger nanoeukaryotes within phytoplanktonic communities^26^. Low nutrient levels and dominance of eukaryotic phytoplankton may favour a higher fraction of the primary productivity allocated to lipids rather than other macromolecules in some regions of the Arctic Ocean compared to other oceans^29,30^. Considering that phytoplankton lipids contribute significantly to the transfer of energy to the Arctic Ocean food web^31^ and that NLs can account for up to 80 % of the lipid pool of the Arctic Ocean phytoplankton^32,33^, phytoplankton NLs may be of major importance for the Arctic Ocean food web.

The metabolism of NLs, mainly PHAs, also play important ecological functions for prokaryotes. PHAs as a storage NL help to store carbon reserve when exposed to nutrients imbalance (N, P, S limited and abundant C), environmental stress (temperature fluctuations, osmotic changes) or feast-famine cycles^34,35^. But PHAs have other important functions: as a reduced carbon form, it helps mitigate the effect of reactive oxygen species due to cold and oxidative stress^36,37^ or protects the cell against ultraviolet radiation^38^. In the ocean, PHA-producing bacteria have been described and isolated from oil-degrading^39^ and plastic-degrading communities^40^ and are generally taxonomically diverse^41^. PHA biosynthesis has also been identified in cold oceanic environments, such as the Antarctic Ocean^42,43^ and the Baltic Sea^44^. The metabolic activity of ocean microbial communities during the Arctic polar night shows that prokaryotes survive and use energy^3^, which implies access to an energy source. In other systems such as soils, it has been shown that storage of energy reserve helps prokaryotes to survive resource-scarce winters^45^. The ability to synthesize PHAs could therefore be very advantageous for the bacteria of the Arctic Ocean to survive through the polar night. However, we possess very scarce information on the capacity and diversity of Arctic Ocean prokaryotes to produce and degrade PHAs. In addition, NLs produced by phytoplankton could be consumed by microbial heterotroph populations, representing an important yet unknown fraction of the C transferred within the microbial loop. Despite a possible important ecological role for the microbiomes and the C cycle of the Arctic Ocean, NL metabolism, its phylogenetic diversity and cycling is poorly understood.

In this study, we postulated that NL production would be more prevalent in phytoplankton of the Arctic Ocean compared to other oceans and that NLs are important carbon and energy sources for heterotrophic prokaryotes. We also investigated if the prokaryotes produce NLs and what C and energy sources they use to feed NL production. We used metagenomics to explore NL metabolic genes in the picoeukaryotic and prokaryotic communities of the Arctic Ocean and compared with other world oceans, as well as determined the taxonomic diversity of microbial species involved. We then determined if heterotrophic prokaryotes would be able to consume NL from phytoplankton origin. And we finally established a link between ecologically important carbon sources such as terrestrially-derived aromatic compounds and NL production in prokaryotes. This work sheds a new light on NL metabolism in Arctic Ocean microbiomes as an important contributor to the Arctic Ocean biogeochemical cycles.

## Materials and Methods

### Sampling, DNA and RNA extraction

Samples were collected in September 2017 during the Joint Ocean Ice Study cruise to the Canada Basin. We analyzed 22 metagenomes generated from samples collected across the water column of the Canada Basin (**Table A1**). Throughout the water column, 8 depths corresponding to specific water masses were sampled (**Table A1**): the surface (5 m and 20 m depth) characterized by fresher water due to riverine input and ice melt, the subsurface chlorophyll maximum (SCM), 2 samples in the fluorescent dissolved organic matter maximum, in the halocline (FDOMmax: at salinity of 32.3 and 33.1 PSU, referred as 32.3 and 33.1), and deeper water from Atlantic origin at the temperature maximum (referred as Tmax), 1000 m depth (Atlantic water, further referred as AW) and 10 or 100 m above the bottom (further referred as bottom).

We filtered 14L of seawater for each sample sequentially through a 3 μm pore size polycarbonate track etch membrane filter (AMD manufacturing, ON, Canada) and a 0.22 μm pore size Sterivex filter (Millipore, MA, USA). Filters were stored in RNALater (ThermoFischer, MA, USA), and kept frozen at −80^0^C until processing in the lab. DNA was extracted following the method described in^97^. Briefly, the preservation solution was expelled and replaced by a SDS solution (0.1 M Tris-HCl pH 7.5, 5% glycerol, 10 mM EDTA, 1% Sodium Dodecyl Sulfate) and incubated at room temperature for 10 min and then at 95^0^C for 15 min. The cell lysate was then ultracentrifuged at 3,270 x g. Proteins were precipitated with the protein precipitation solution MCP (Lucigen, WI, USA), and supernatant was collected after centrifugation at 17,000 x g for 10 min at 4^0^C. DNA was then precipitated with 0.95 volume of isopropanol and rinsed twice with 750 μL ethanol before being air dried. The DNA was resuspended in 25 μL of low TE buffer, pH8 (10 mM Tris-HCl, 0.1mM EDTA) and stored at −80^0^C.

### Metagenomic sequencing, assembly, and annotation

Sequencing, assembly, and annotation were performed by the Joint Genome Institute (CA, USA). Metagenomes were sequenced on the Illumina NovaSeq platform, generating paired-end reads of 2×150 bp for all libraries. Single assemblies were created by JGI for each individual sample using SPAdes^98^ with kmer sizes of 33, 55, 77, 99, 127 bp. Gene prediction and annotation was performed using the DOE Joint Genome Institute Integrated Microbial Genomes Annotation Pipeline v.4.16.5^99^.

### Computation of gene abundance, expression, and transcripts abundance profiles

Metagenomics data files containing genes IDs, gene annotations, gene depth of coverage and other gene information were retrieved from the DOE Joint Genome Institute Integrated Microbial Genomes (JGI/IMG, https://img.jgi.doe.gov) repository (**Table A1**). Metagenomes from the Canada Basin originate from samples collected by our lab. Metagenomes from other world oceans originate from samples collected by others and annotated by JGI/IMG (**Table A3**). The abundance of a KEGG ortholog number (KO, gene family) in a metagenome was calculated by summing the depth of coverage of all genes annotated with this KO. KO abundance matrices therefore represent the metagenomic profiles across the samples. As samples vary in terms of depth of sequencing (or library size), we normalized the metagenomic profiles to the relative cell number to get a per-cell number of copies using the approach detailed in Salazar *et al*^100^. Specifically, the KO abundances were divided by the median abundance of 10 universal single-copy phylogenetic marker genes (K06942, K01889, K01887, K01875, K01883, K01869, K01873, K01409, K03106, and K03110). These normalized KO abundances can therefore be interpreted as the per-cell number of gene copies for a given protein family (KO).

### Selection of NL storage and degradation pathways and marker genes

Metabolic pathways for the biosynthesis of TAGs, WEs, SEs and PHAs as well as metabolic pathways for the degradation of TAGs, SEs and PHAs were retrieved from MetaCyc (https://metacyc.org, **Table A2**). To evaluate the abundance of various pathways, we used the abundance (see above for abundance calculation) of genes coding for various protein families (KO) that catalyze key reactions (EC) in the NL biosynthesis and degradation pathways. The key reactions for TAGs (EC:2.3.1.20 – diacylglycerol O-acyltransferase/wax synthase and EC:2.3.1.158 – phospholipid:diacylglycerol acyltransferase), WEs (EC:2.3.1.75 – wax synthase) and SEs (EC:2.3.1.26 – sterol O-acyltransferase) biosynthesis were selected as the esterification reaction between a FA and diacylglycerol, a fatty alcohol and sterol respectively. The marker enzyme for the degradation of TAGs (EC:3.1.1.3 – TAG acylhydrolase) and SEs (EC:3.1.1.13 – SE esterase) was responsible for the cleavage of the ester bond to free a FA from TAGs and SEs respectively. The key reaction chosen for the synthesis of PHAs (EC:2.3.1.304 – PHA synthase) was the esterification involved in the elongation of polyhydroxyalkanoate while the de-esterification (EC:3.1.1.75 – PHA hydrolase) was chosen as the key reaction for the PHA degradation pathway. As EC:2.3.1.304 was created in 2021, after the annotation of our metagenomes in 2018, we could not find any genes annotated with EC:2.3.1.304 in our dataset. Instead, we used the KEGG ortholog annotation K03821. There was no metabolic pathway for the degradation of WE in Metacyc. We therefore selected enzymes previously reported to cleave the ester bond of wax ester as marker enzymes: wax ester hydrolase (EC:3.1.1.50) and cutinase (EC:3.1.1.74).

### Selection of genes involved in exogenous fatty acid transport

To retrieve genes involved in the import of exogenous FAs in prokaryotes, we surveyed the literature of exogenous FA use by prokaryotes^46,47,67,103–105^ (**Table A2**). Based on the literature survey, we retrieve genes belonging to the well-known *fadL/fadD/fadR* (K06076/K01897/K13770) system of long-exogenous FA import as well as the short-chain FA transporter *fadK* (K12507). We also retrieved genes for import system that were non-specific for exogenous FAs, but have been reported to facilitate the import of exogenous FAs: the porin *ompF* (K09476), permease *mce1* (K02066) and an ABC transported (K24820). We also retrieved genes that we implicated in the funneling of exogenous FA to phospholipids: the *fakA/fakB* (K07030/K25232) and the *aas* gene (K05939)

### Taxonomic assignment of neutral lipid metabolism marker genes

To assign a taxonomy to genes annotated to marker reactions of neutral lipid metabolic pathways, we first grouped genes per water column and used CD-hit^106^ to dereplicate the genes at 95% identity. We searched the dereplicated set of genes against the NCBI nr database (download on August 27^th^ 2021) using DIAMOND blastp^107^. The DIAMOND output was imported to MEGAN^107^ using the January 2021 mapping file (“megan-map-Jan2021.db”). In MEGAN, we set the lower common ancestors’ parameters at a minimum e-value of 1×10^-20^ and a top precent of 1%. We exported the file containing the taxonomic affiliation of the genes and processed it with a custom-made *R* code.

### Binning

Single sample metagenomic assemblies and metagenomic co-assemblies of samples from the Canada Basin and Amundsen Gulf (**Table A7**) were performed to recover a greater taxonomic diversity of metagenome assembled genomes (MAGs). Co-assemblies of 24 samples from the Canada Basin, 11 samples from Canada Basin surface, 7 samples from the Amundsen Gulf, and individual assemblies of all 31 samples from the Canada Basin and Amundsen Gulf were performed. All metagenomic assemblies were generated using MEGAHIT (v.1.2.7)^108^ with k-mer sizes of 27,37,47,57,67,77,87. The input metagenome reads for each assembly were mapped to the assembly with BWA (v.0.7.17)^109^ using the mem option. The mapping results were processed using jgi_summarize_bam_contig_depths from MetaBAT2 v.2.12.1^110^ and metagenomic binning was performed for each of the assemblies using MetaBAT2 (v.2.12.1) with default settings, resulting in 4824 genome bins. Genome quality was evaluated using CheckM v.1.0.114^111^ with the lineage_wf workflow. The 924 bins with >50% completeness and <10% contamination and strain heterogeneity, were considered at least medium quality MAGs^112^. The MAGs with >50% completeness and <10% contamination and strain heterogeneity were dereplicated with dRep^113^ using 95% ANI cut-off to remove species level redundancy, resulting in 664 representative MAGs. Dereplicated MAGs with >50% completeness and <10% contamination and strain heterogeneity were taxonomically classified with the GTDB-tk (v.1.3.0)^114^ using the classify_wf workflow.

### Annotation of MAGs

We used Prokka (v.1.12)^115^, implementing Prodigal (v.2.6.3)^116^, to predict and annotate genes in MAGs. INFERNAL (v.1.1.2)^117^ was used to predict ribosomal RNA genes against Rfam (v.14.2)^118^, KEGG, KofamScan/KofamKOAL^119^ and Metacyc, using default settings and a bitscore-to-threshold ratio of 0.7.

### Metabolic reconstruction in MAGs

The presence of marker genes involved in the metabolism of neutral lipids were retrieved based on their annotation with EC and KO numbers associated with markers reactions (see above). The number of genes coding for enzymes that assemble, modify and breakdown oligo- and polysaccharides in MAGs were retrieved using genes annotated with EC numbers corresponding to carbohydrate-active enzymes (CAZymes, **TableA4**) as described in^120^. The number of genes coding for enzymes involved in AC degradation (**Table A5**) were retrieved in MAGs using genes annotated with dioxygenases (EC:1.13.11.-) and monooxygenases (EC:1.14.13.-) from aromatic compound degradation pathways within Metacyc (https://metacyc.org/). The number of genes annotated with transporters for carbohydrates, proteins/aa and lipids were retrived in MAGs based on their KO annotation from the KEGG database (https://www.genome.jp/kegg/, **Table A7**)

### Statistical tests

To calculate the co-occurrence between NL metabolism marker genes and FA transport genes in MAGs, we computed the Sorensen index using the presence and absence of these genes in MAGs. The p-value were calculated using 9999 permutations. P-values < 0.05 were considered significant.

## Results

### Physicochemical structure of the Canada Basin water column

This study focused on the microbiomes of the Canada Basin sampled along a latitudinal transect from 73 ^0^N to 81^0^N. We collected samples during the summer and fall (September) through the water column of the salinity-stratified Canada Basin, targeting four distinct water column features (**Figure 1a-b, A1, Table A1**). The surface (5 m and 20 m depth) and subsurface chlorophyll maximum (SCM; 55-95 m) are within the photic zone. The surface samples were characterized by lower salinity (25-29 PSU) and high phytoplankton cell abundance, while the SCM had a high level of fluorescent dissolved organic matter (FDOM) and the highest bacterial cell abundance (**Figure 1a-b, A1**). Below the photic zone, we retrieved samples from Pacific-origin water, at salinities of 32.3 and 33.1 PSU. These waters were characterized by the highest levels of FDOM, representing the FDOMmax feature (**Figure 1a-b, A1**), where organic matter of terrestrial origin accumulates. The phytoplankton abundance sharply decreased close to zero in the FDOMmax, while it contained the highest nutrient (nitrates, phosphates, silicates) concentrations of the water column. The water below 400 m depth is of Atlantic origin. In this water, referred to as deep water in this study, we collected samples at the temperature maximum (Tmax), at 1000 m depth and at the bottom. Deep samples were characterized by the highest salinity and nutrient concentrations.

**Figure 1:**
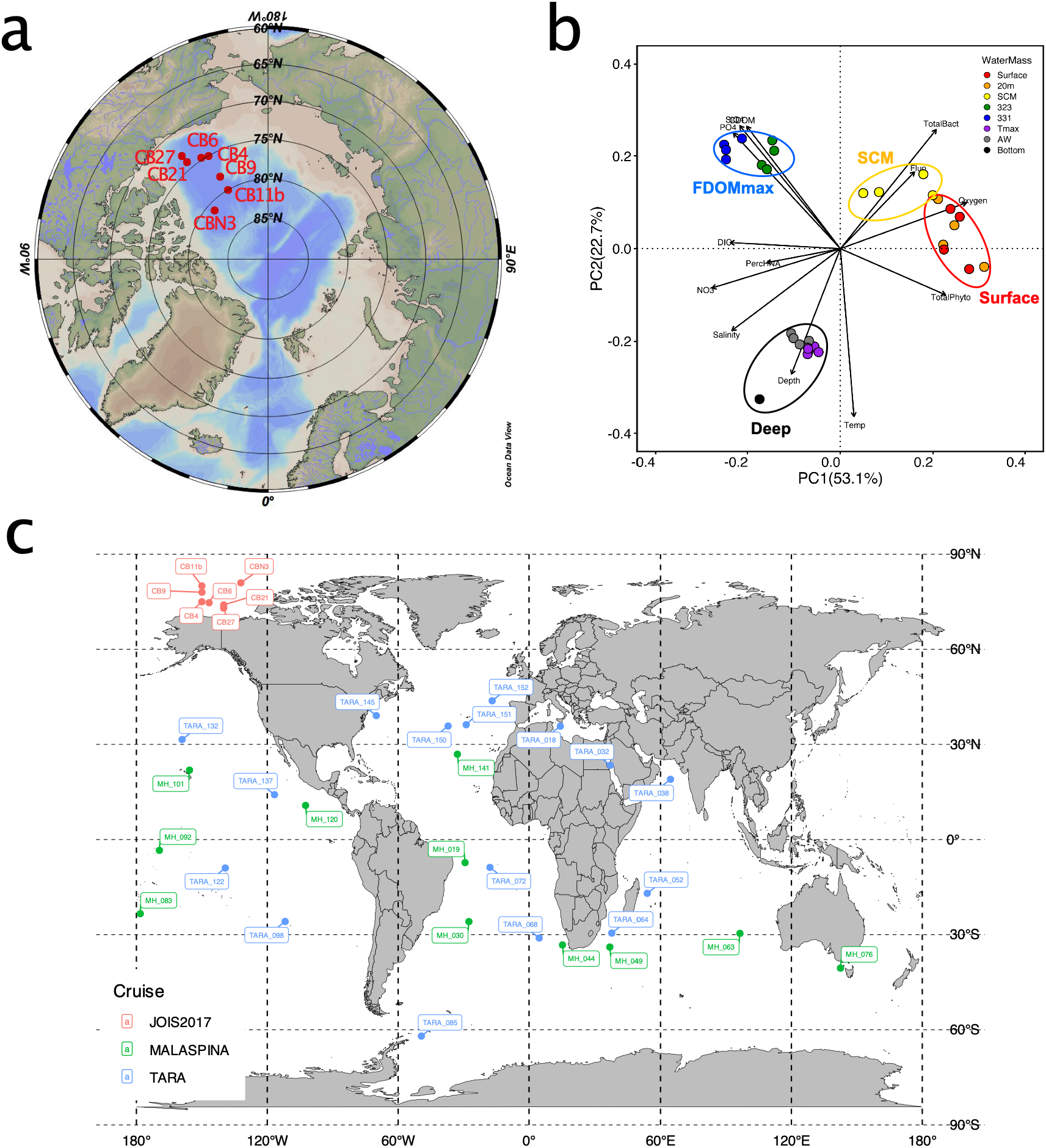
Selection of the water column features sampled in the Canada Basin of the Arctic Ocean. a) Map of the station sampled in the Canada Basin. b) Principal component analysis of the samples based on their physicochemical parameters that show the differences among the four water column features: the surface (red circle), the SCM (yellow circle), the FDOMmax (blue circle) and the deep samples (black circles). SCM= subsurface chlorophyll maximum; FDOMmax= Fluorescent dissolved organic matter maximum; 323=salinity of 32.3 PSU; 331= salinity of 33.1 PSU; Tmax= temperature maximum; AW= Atlantic water. c) Map of the stations in the global ocean from which metagenomes were retrieved. Red=Samples retrieved from the Canada Basin for this study, blue and green= samples from publicly available metagenomes from the TARA (blue) and Malaspina (green) expeditions.

### Biosynthesis and degradation of neutral lipids in Arctic and global ocean microbiomes

We first sought to examine the biogeography of NL metabolic genes in Arctic Ocean microbiomes and how it compared to other regions of the global ocean. We therefore retrieved genes involved in the NL biosynthesis and degradation pathways from metagenomes (**Figure A2, A3**). We also sought to determine the taxonomic diversity of NL metabolic genes within the Arctic Ocean microbiomes. All the NL biosynthesis pathways were complete in the photic zone (surface and SCM) and FDOMmax of the Arctic Ocean (**Figure A4, left panel**). All the NL degradation pathways were complete in the photic zone and the FDOMmax of the Arctic Ocean, except the TAG degradation pathway that was complete only in the surface (**Figure A4, right panel**).

To quantify the biogeographic distribution of NL metabolism in the ocean microbiomes we focused on the abundance of gene families (KEGG Orthologs) catalyzing key steps of NL biosynthesis and degradation (**Figure 2a-c, A2, A3, Table A3)** and normalized by the median abundance of 10 conserved single-copy gene families (see methods). The diacylglycerol acyltransferase (DGAT), wax ester synthase (WS) and steryl ester synthase (SES) catalyze the esterification of a fatty-acyl moiety with diacylglycerol, a fatty-alcohol, or a sterol, respectively, the key step in these NLs biosynthesis pathways. The deesterification of these three NLs to free a fatty-acyl is catalyzed by either the TAG lipase, WE lipase or SE lipase^22^. The key steps in PHA metabolism are the polymerization and depolymerization steps, catalyzed by the PHA polymerase and PHA depolymerase, respectively^14^.

**Figure 2:**
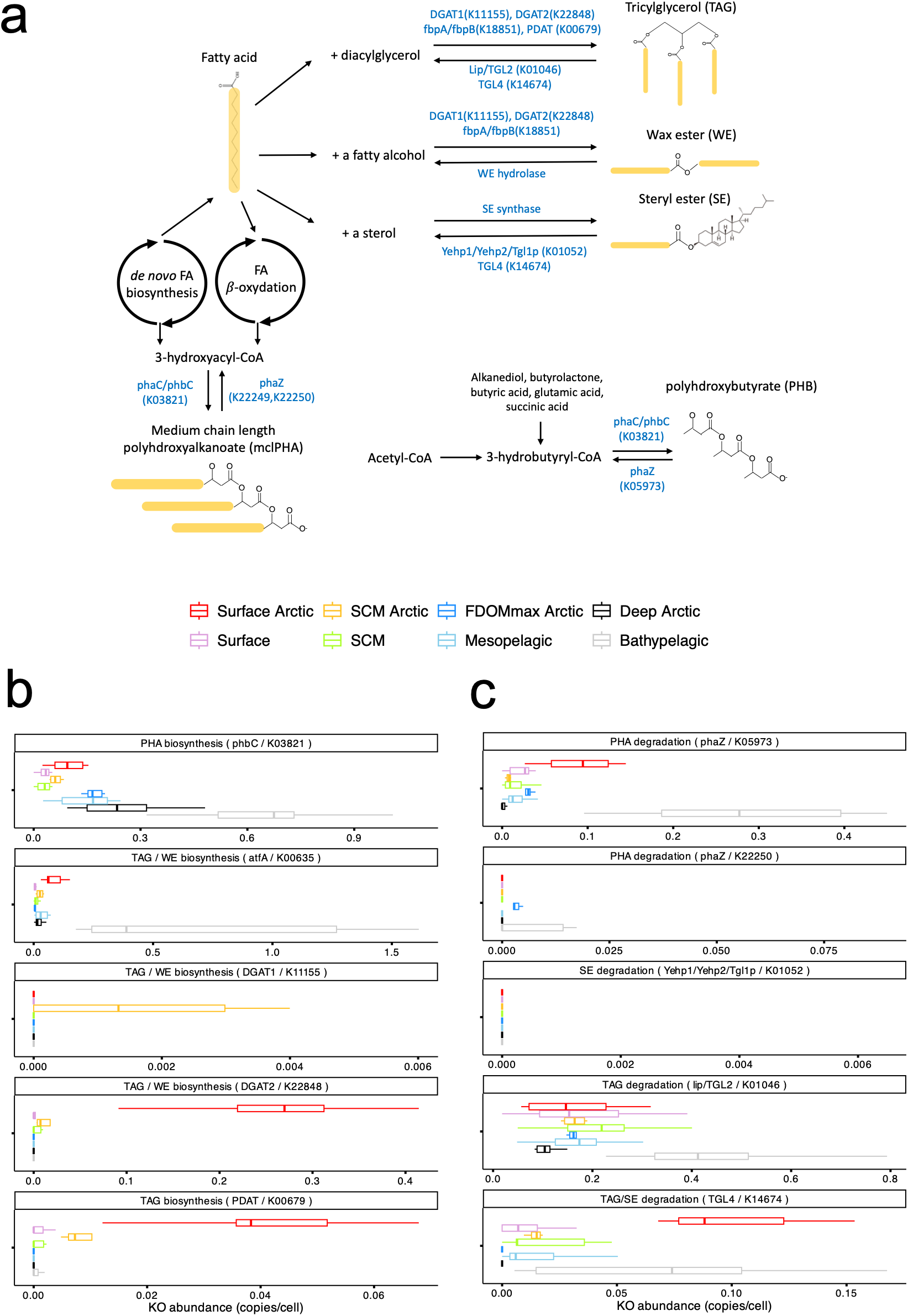
Metabolism of neutral lipids in the Canada Basin and the global ocean microbiomes. a) Schematics of biosynthetic pathways used to produce the neutral lipids triacylglycerols, wax esters, steryl esters and polyhydroxyalkanoates. b) Abundance of genes coding for protein families that catalyze key steps of the neutral lipids biosynthetic pathways within metagenomes of the Canada Basin (red, yellow, blue and black) and the global ocean (shades of grey).

TAG biosynthesis genes were generally more abundant in the photic zone of the Arctic Ocean compared to the photic zone of other oceans (**Figure 2b)**. This trend was particularly marked for genes with ws/dgat activity (DGAT2/K22848, up to 0.28 copies/cell) and with phospholipid:diacylglycerol transferase (pdat) activity (PDAT/K00679, up to 0.039 copies/cell), with abundances more than 10 times higher in the photic zone of the Arctic ocean microbiomes compared to the microbiomes of other oceans. In the Arctic Ocean, all genes within these two families were associated with eukaryotic phytoplankton groups in the *Mamelliales* order, including *Micromonas* and *Bathycoccus* species and the *Pelagomonadales* order (**Figure 3**). We also identified one gene family with ws/dgat activity (WS/DGAT - K00635) that displayed prokaryotic origin, within *Gammaproteobacteria*.

**Figure 3:**
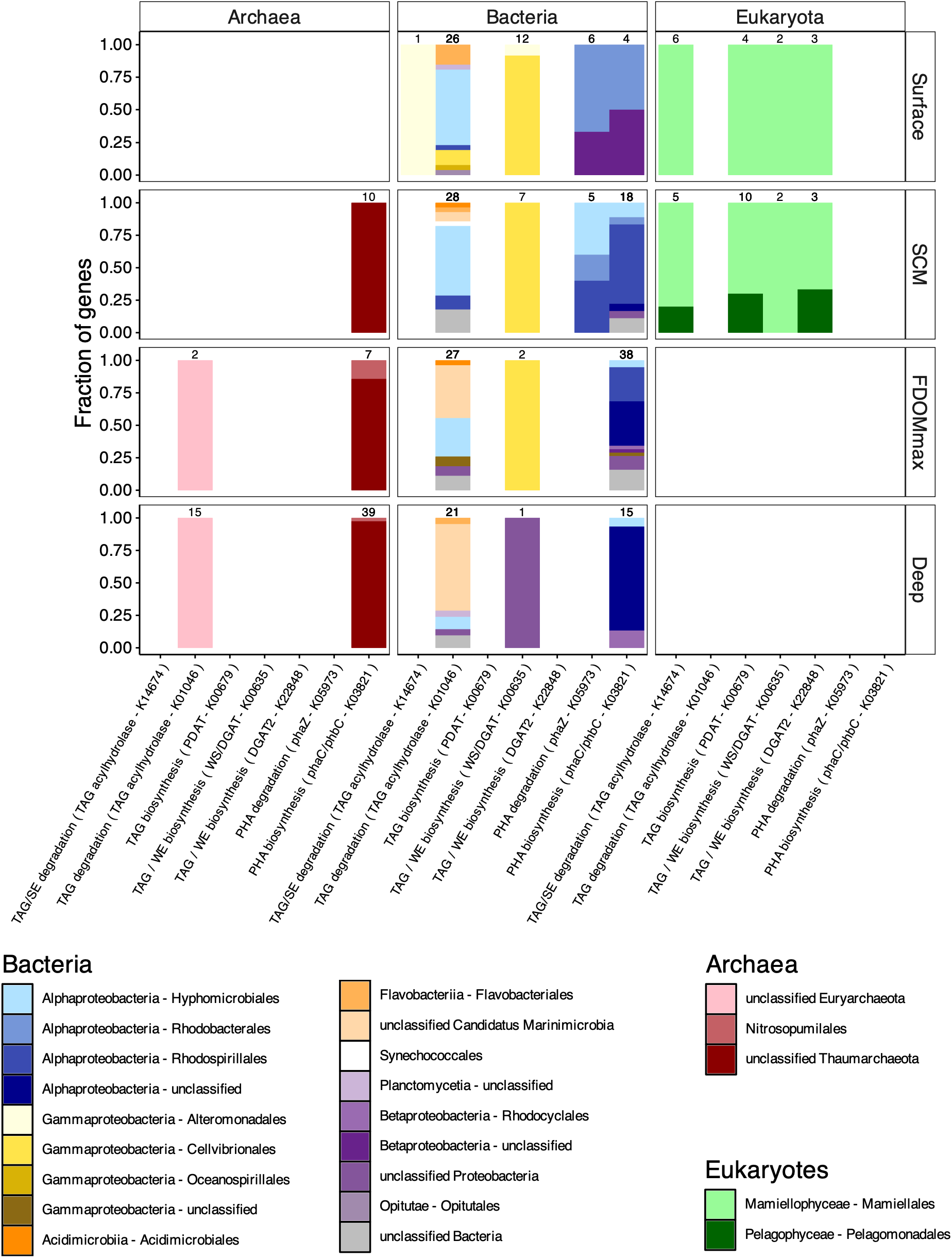
Taxonomic affiliation of genes coding for protein families that catalyze key step of neutral lipid biosynthetic and degradation pathways. The taxonomy is separated by domain and water column features of the Canada Basin. The numbers at the top of each bar represent the total number of gene clusters (clustered at 95% identity) obtained for a protein family (KO number) in a water column feature.

The TAG degradation lipase TGL4 genes (K14674) were ∼10 times more abundant in the Arctic Ocean’s photic zone microbiomes (0.85 copies/cell, **Figure 2b**) than in other ocean photic zone microbiomes. Similarly to the PDAT gene, the TGL4 gene was of eukaryotic phytoplankton origin, with only one variant from *Gammaproteobacteria* (**Figure 3**). The second TAG degradation lipase (*lip*/TGL2 - K01046) we found was exclusively of prokaryotic origin, with a different and more diverse taxonomic distribution than the WS/DGAT gene family K00635 (**Figure 3**). Interestingly the abundance of the TAG degradation lipase *lip*/TGL2 (0.18 copies/cell) was higher than the TAG synthesis gene WS/DGAT (K00635, 0.05 copies/cell) within the Arctic Ocean surface. These results show that in the Arctic Ocean photic zone, the main TAG producers are the eukaryotic phytoplankton, and that, within the prokaryotic communities, the TAG degraders are different, more diverse and more abundant than the TAG producers. That might suggest the existence of a “lipotrophic” prokaryotic community, degrading and using TAGs produced by the eukaryotic phytoplankton fraction in the Arctic Ocean surface.

The abundance of PHA biosynthesis genes generally increased with depth (**Figure 2b**). Both the PHA biosynthesis (*phaC*/*phbC*) and degradation (*phaZ*) genes were more abundant in the Arctic surface microbiomes (0.09 copies/cell) than in the surface of other oceans’ microbiomes (0.03 copies/cell) but less abundant in the Arctic Ocean deep waters (0.25 and 0.01 copies/cell, respectively) than the bathypelagic waters of other oceans (0.67 and 0.28 copies/cell, respectively) (**Figure 2b**). In the Arctic Ocean both PHA biosynthesis and degradation genes were exclusively from prokaryotic origin, with similar taxonomic distribution within *Alphaproteobacteria* and *Betaproteobacteria*.

### Fatty acid import in Arctic and global ocean microbiomes

The existence of a “lipotrophic” group of bacteria in the photic zone of the Arctic Ocean would necessitate genes involved in exogeneous fatty acid (FA) import across the cell membrane. Indeed, exogeneous FAs are freed from NLs outside of the cell, and then imported to be used as an energy source or other purposes. We therefore asked if the microbial communities of the Arctic Ocean would be enriched in FA transport genes (**Figure 4a**). We also asked how the imported exogeneous FA would preferentially be used: as an energy source or incorporated in phospholipids. We therefore explored the distribution of exogeneous FAs import and use genes in the Arctic Ocean and global ocean microbiomes.

**Figure 4:**
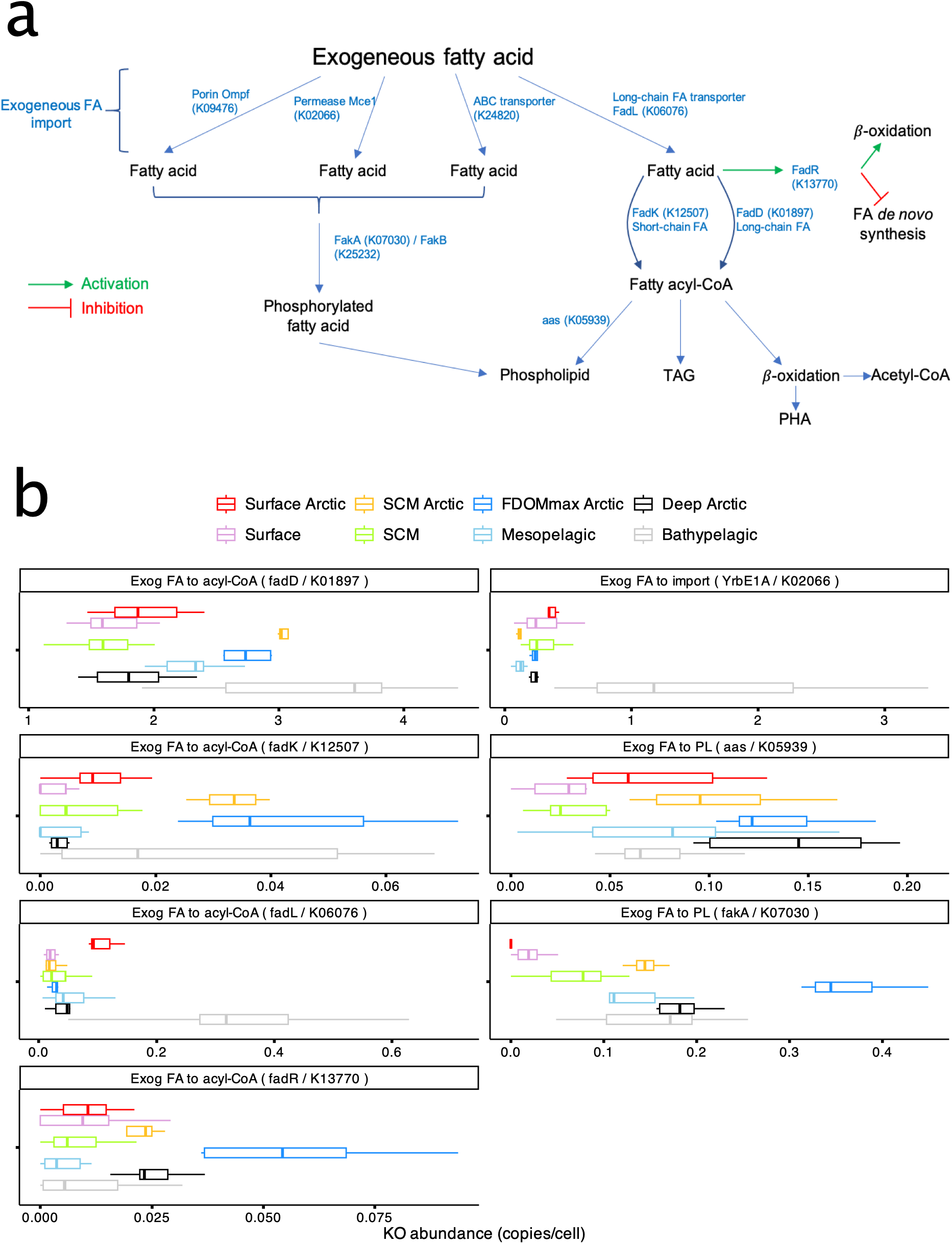
Import of exogenous fatty acids (FAs) in the Canada Basin and the global ocean microbiomes. a) Schematics of various transporters that have been reported to selectively import FAs or facilitate the import of FA non-selectively. b) Abundance of genes coding for protein families involved in exogenous FA import within metagenomes of the Canada Basin.

The transmembrane import system for FAs *fadL/fadD/fadR*, often found on an operon is a widely studied system that directs long-chain FAs to β-oxidation^46,47^: the *fadL* transporter is specific to long-chain FAs and coupled to the FA acyl-CoA synthase *fadD* which catalyzes the acetylation of FA that is the substrate of β-oxidation. The *fadR* gene, a transcription factor, completes the system by activating the β-oxidation pathway and suppressing *de novo* FA synthesis. The abundance of *fadL* was significantly higher in the photic zone of the Arctic Ocean (**Figure 4b**), especially at the surface (0.1 copies/cell), than in the photic zone of the rest of the global ocean (<0.01 copies/cell). Similarly to the abundance, the taxonomic diversity of the *fadL* gene was also highest in the photic zone (surface and SCM), and decreased drastically in the aphotic zone, suggesting a connection to the eukaryotic phytoplankton TAG producers (**Figure 2b**). This supports the hypothesis that a fraction of the microbial community of the surface of the Arctic Ocean may use exogeneous FAs from phytoplankton-produced TAGs. The *fadL* genes were assigned to *Gammaproteobacteria* and *Bacteroidetes*.

As an alternative to β-oxidation, exogeneous FAs can also be incorporated in cellular phospholipids. Considering the energy requirement of *de novo* FA biosynthesis, this potentially saves a considerable amount of energy for the cell. We detected two genes involved in incorporating exogeneous FA into phospholipids (**Figure 4a**): the *aas* gene that transfers the fatty acyl moiety from the acyl-[acyl-carrier] protein to the phospholipid (PL)^48^ and the *fakA* gene that phosphorylates a FA, a step necessary for the incorporation of FA in PLs^49^. The abundance of the *aas* gene peaked in the FDOMmax (0.125 copies/cell) and was higher in all the Arctic Ocean water masses than corresponding water masses outside of the Arctic Ocean (**Figure 4b**). Similarly, the taxonomic diversity of the *aas gene* peaked in the FDOMmax where it was mostly assigned to *Verrucomicrobia* (**Figure S5**). The *fakA* gene displayed an even higher abundance (0.34 copies/cell in the Arctic FDOMmax), where it was taxonomically assigned to *Chloroflexi, Marnimicrobia* and *Acidimicrobiia* (**Figure A5**).

### Identification of NL metabolic genes in metagenome-assembled genomes

Based on our assumption that FAs derived from NLs produced by eukaryotic picophytoplankton can be used by a fraction of the heterotrophic prokaryotic communities – the “lipotrophs” - we sought to determine the taxonomic diversity of this lipotrophic fraction of the prokaryote community. We also asked whether the imported FAs could be used as a source of energy or incorporated within NLs or PLs. To that end we reconstructed metagenome-assembled genomes (MAGs) from our metagenomic data to determine the presence and co-occurrence of exogeneous FAs import genes and NL metabolism genes. After filtering for genomes greater than 50% completeness and less than 10% contamination and strain heterogeneity, we obtained a set of 664 MAGs.

We observed a clear taxonomic difference for the composition of FAs import genes between MAGs involved in NL metabolism and those that are not: at the exception of MAGs in the *Actinobacteria* phylum, the vast majority of MAGs possessing NL metabolism genes did not possess the *fakA* gene. Conversely, MAGs without any NL metabolism genes almost all possessed the *fakA* gene that funnels exogeneous FAs to phospholipids and were in the *Chloroflexota* phylum (**Figure 5, A6**). These observations suggest that taxa in the *Chloroflexota* phylum use exogeneous FAs mainly for phospholipids, saving the cost of *de novo* FA synthesis.

**Figure 5:**
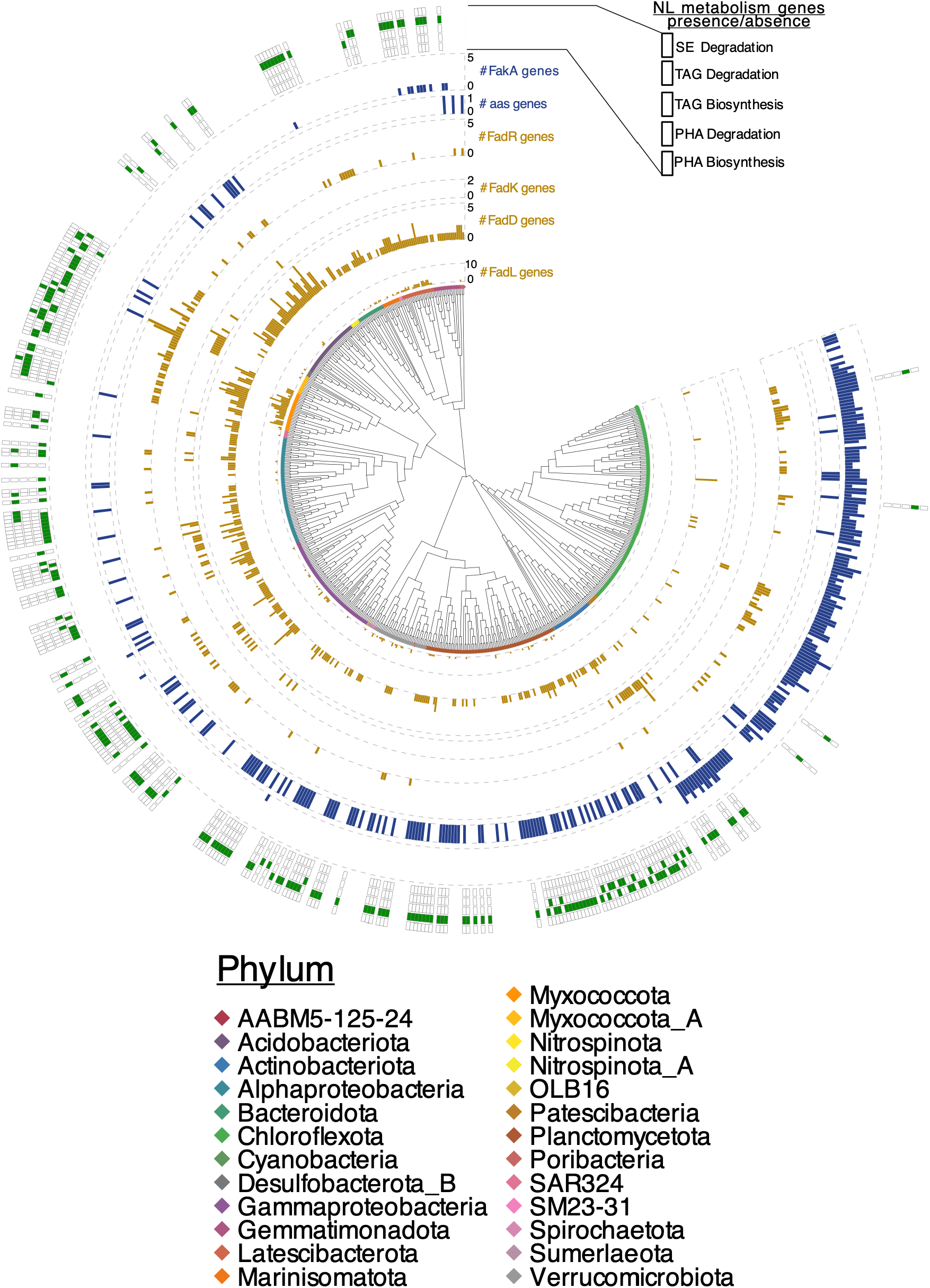
Phylogenetic tree of bacterial metagenome assembled genomes (MAGs) displaying neutral lipid metabolism and exogenous FA import genes. The presence of gene coding for protein families catalyzing key steps of NL biosynthesis and degradation in a MAG is represented by a green cell. The number of genes involved in the import of FA and funneled to β-oxidation (dark yellow) or to membrane lipids (blue) in each MAG is represented by a vertical bar.

Within MAGs possessing NL metabolism genes, a significant fraction (22%, 149 MAGs) in various phyla (*Gammaproteobacteria, Verrucomicrobiota, Planctomycetota, Gemmatimonadota, Poribacteria, Marinisomatota*) possessed a TAG degradation gene but no TAG biosynthesis gene (**Figure 5**). The majority of those MAGs also possessed the *fadL* and *fadD* genes for the transport and activation of long-chain exogeneous FAs with CoA as evidenced by the co-occurrence of *fadL* and *fadD* with TAG degradation genes (**Figure A6**). These results reinforce our hypothesis that NL derived from eukaryotic picophytoplankton can serve as an important energy source for a significant fraction of the heterotrophic prokaryotic community. A small fraction of the MAGs (7%, 46 MAGs) was able to synthesize TAGs as they possessed the TAG synthesis gene (**Figure 5**). These MAGs were found in the phyla *Myxoccocota, Actinobacteria* and *Planctomycetota* and a large fraction also possessed the *fadL* gene (**Figure 5, S6**), suggesting that they might incorporate exogeneous FAs into NLs. Interestingly most MAGs in the *Myxococcota* phylum possessed many copies of the *fadL* gene, highlighting the possible importance of exogeneous FAs for TAG biosynthesis in these taxa. The *phaZ* gene, involved in PHA degradation was found in only 2% of the MAGs (10, **Figure 5**). A larger fraction of MAGs (6%, 43 MAGs) possessed the *phaC* and *phbC* genes involved in PHA biosynthesis. These MAGs were mainly in the *Alphaproteobacteria and Gammaproteobacteria* phyla, which differed from the taxonomy of TAG biosynthesis MAGs. The MAGs with PHA biosynthesis capacity generally had very few or no *fadL* gene, suggesting that exogeneous FAs were not used to feed PHA biosynthesis (**Figure A6**). These results show that exogeneous FAs may be an important source of energy for heterotrophic prokaryotes of the Arctic Ocean but could also support NL accumulation in a small subset of the communities.

### Organic carbon sources for TAG and PHA biosynthesis in Canada Basin microbiomes

Our results highlighted the capacity of bacterial taxa to synthesize both TAG and PHA in the Arctic Ocean. The storage of NLs may help these bacteria to store C reserve in anticipation of the C shortage during the Arctic winter. We also found that taxa able to synthesize TAG were generally able to import FA while those able to synthesize PHA were not. We therefore asked if other sources of carbon with ecological importance in the Arctic Ocean could be used by bacterial taxa to potentially feed NL biosynthesis. Studies investigating the storage of NLs in model species of prokaryotes have shown that they can use sugars^50^ or aromatic compounds^51^ as both a C and energy source to feed NL accumulation. Sugars and polysaccharides are commonly produced by phytoplankton and have been shown to be ecologically relevant to sustain bacterial communities^52^. Recently our group demonstrated that the unique capacity of Arctic Ocean microbiomes to degrade aromatic compounds^53^ is an evolutionary adaptation to use the disproportionately high amount of organic matter from terrestrial origin in the Arctic Ocean. We therefore investigated the capacity of MAGs involved in NL biosynthesis to use various carbon sources to feed NL biosynthesis.

We found that MAGs with PHA and TAG biosynthetic genes were phylogenetically different, separated in the water column and seemed to favored different carbon sources. MAGs with PHA biosynthetic capacity, in the *Alphaproteobacteria* and *Gammaproteobacteria* phyla, were generally found in the SCM and FDOMmax (**Figure 6**), where we find an accumulation of organic matter from terrestrial sources and hence aromatic compounds^54^. These MAGs possessed on average a higher number of aromatic compounds degradation genes than the MAGs with TAG biosynthetic capacity (**Figure A7**). Conversely, MAGs with TAG biosynthetic capacity belonged mainly to the *Myxococcota* and *Planctomycetota* phyla, were preferentially found in deep waters (**Figure 6**), and possessed a higher number of genes coding for carbohydrate transporters and carbohydrate active enzymes (CAZymes) (**Figure A7**) than PHA biosynthetic MAGs, indicative of a preference for carbohydrates as a carbon source. Within the TAG biosynthetic MAGs, *Actinobacteria* was the only phyla found in the upper water column (SCM and FDOMmax), and similarly to other MAGs in SCM and FDOMmax with PHA biosynthesis capacity, also possessed a higher number of aromatic compound degradation genes, as well as genes for amino acids/peptides transporters (**Figure 6**).

**Figure 6:**
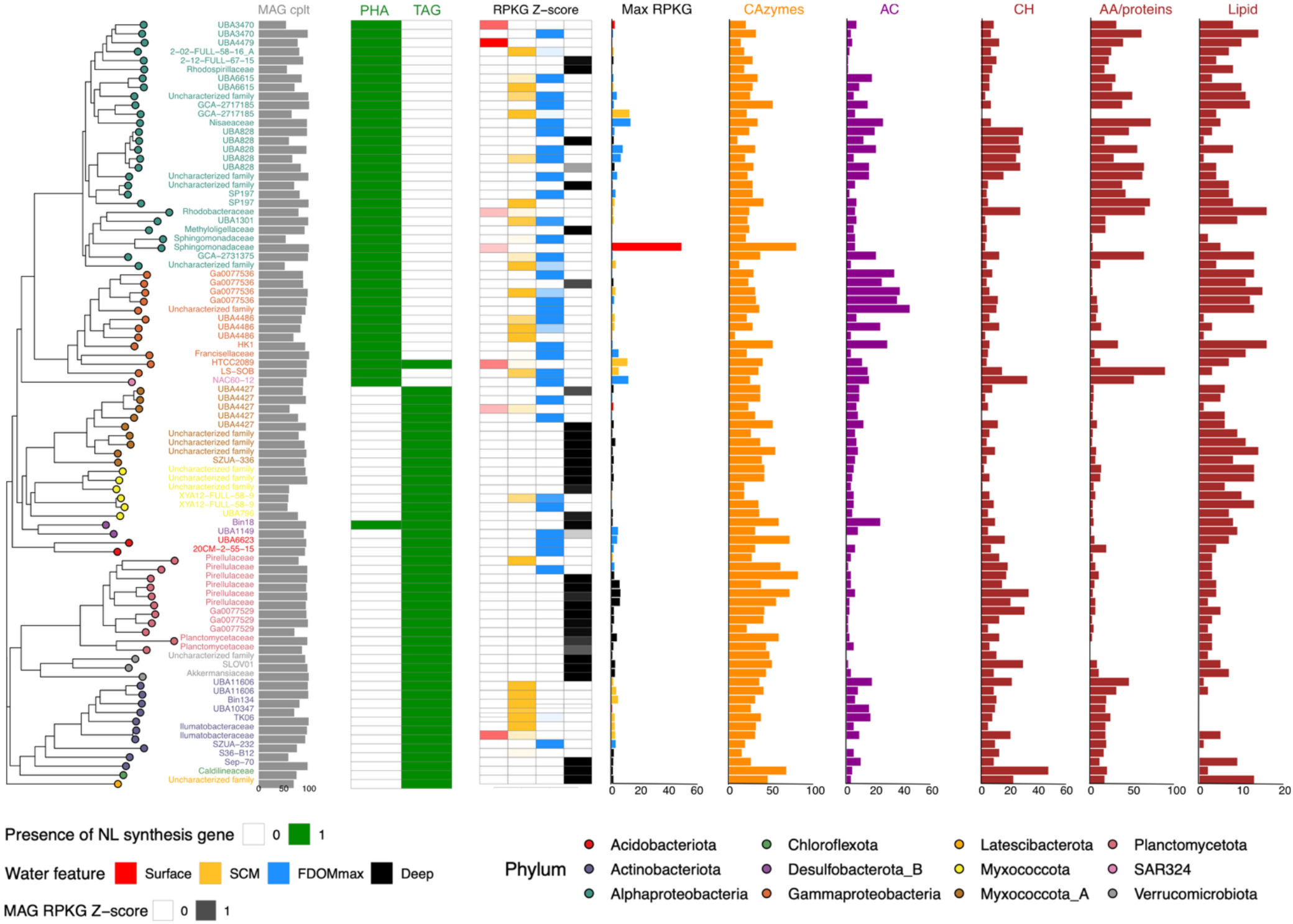
Phylogenetic tree of bacterial metagenome assembled genomes (MAGs) displaying genomic capacity for NL synthesis and use of various carbon sources. Grey bars represent the MAG completeness (%). Green cells represent the presence of genes coding for the protein families catalyzing key steps of triacylglycerol or polyhydroxyalkanoate cycling. The red/yellow/blue/black heatmap represents the z-score of a MAG abundance over all the samples, averaged for each water feature. The red/yellow/blue/black bar plot represents the maximum abundance of each MAG, color-coded by the water feature of the samples in which the MAG was the most abundant (red=Surface, yellow= SCM, blue= FDOMmax, black=deep). The orange, purple and dark red bar plot represent, for each MAG, the number of genes coding respectively for CAZymes (orange), AC degradation enzymes (purple) and carbohydrates (CH), amino acids/proteins (AA/proteins) and lipid transporters (dark red).

## Discussion

### The importance of phototrophic eukaryotes for NL storage in the Arctic Ocean

Our work shows that the metabolism of NLs may be more important than previously recognized for the survival of Arctic phytoplankton, especially through winter. It has been reported that the use of accumulated NLs coupled with reduced metabolism is indeed a strategy for some Arctic phytoplankton to overwinter during the dark polar night^19^. However, our work goes further and demonstrates the capacity for NL biosynthesis in some of the most abundant and ecologically relevant Arctic Ocean phytoplankton taxa, including *Bathycoccus* and *Micromonas.* The advantage provided by NL biosynthesis to *Bathycoccus* highlighted in our work may explain their abundance in the winter prasinophyte taxa^55^. *Micromonas* is ubiquitous in the Arctic Ocean and often dominates phytoplanktonic assemblages^56,57^. The capacity of *Micromonas* to synthesize NLs that we observe in this study may provide them with a significant advantage to dominate phytoplanktonic assemblages across the Arctic Ocean. Beyond these specific taxa, the environmental conditions and seasonal cycles of the Arctic Ocean provide optimal conditions for NL accumulation in phytoplankton. The accumulation of NLs is favored when phytoplankton are faced with an excess of C and energy and a simultaneous limitation in other nutrients (N, P). The cell can then use this C in excess of current metabolic need to store reserves, a process called *surplus storage*. The Arctic summer represents ideal conditions for *surplus storage* in phytoplankton: the low N of summer oligotrophic waters prevents phytoplankton growth, but coupled to long periods of intense light (energy source), constitute optimal conditions for the accumulation of C reserve in the form of NLs^58^. The ability and favorable conditions to store NLs and especially TAGs in phytoplankton could favor eukaryotic phytoplankton over prokaryotic phytoplankton (*Cyanobacteria*) who use glycogen or PHAs as storage compounds ^59^. This could partly explain the absence of *Cyanobacteria* from the photosynthetic communities of the Arctic Ocean^60^.

We expect that the storage of NLs in the Arctic Ocean phytoplankton populations will be enhanced with the changing conditions related to climate change. The Arctic Ocean is freshening, which increases the stratification and prevents nutrients resupply from deep waters ^61^. In addition, climate change observations and current scenarios indicate a longer period of phototrophic activity in the Arctic Ocean due to higher light intensity in the photic zone with sea ice loss^26^. These conditions are bound to further favor the storage of NLs in phytoplankton in the future. Studies have also shown that the taxa capable of NL synthesis highlighted in this work will become more prevalent as climate-change is progressing in the Arctic Ocean ^26^. Given these observations, NL importance in the transfer of energy from phytoplankton to the rest of the food chain could significantly increase in the future.

### TAGs from phytoplankton origin serve as a carbon source for a “lipotrophic” subset of the Arctic Ocean microbiome

We suspect that the higher fraction of lipids in the primary produced C pool of the Arctic Ocean compared to other oceans may be used as an important C source for the heterotrophic microbial populations. Our results show that the diversity of prokaryotic taxa with the capability to degrade TAGs was greater than the diversity of taxa able to synthesize TAGs. This supports the existence of a “lipotrophic” fraction of the microbiome not able to store NLs but able to use exogenous TAGs as C and energy source in the Arctic Ocean. Lipids produced by phytoplankton, either as TAGs or FAs can be released in the water column through various mechanisms such as exudation^62^, viral lysis^63,64^, autolysis^28^ or algicidal bacteria^65,66^. NLs cannot be imported inside the cell, so the TAG acylhydrolase is secreted extracellularly^67^ and the FAs freed from TAG are imported into the cell. The capacity for the “lipotrophic” prokaryotes to use exogeneous TAGs would need the co-existence of TAGs and FA import genes. This is what we observe in our results with the co-occurrence of the TAG hydrolase and genes involved in exogenous FA import in our MAG dataset. This result is important as it reveals a new importance of NL metabolism in the C cycle of the Arctic Ocean.

### Expanded taxonomic diversity of prokaryotic TAG and PHA storers in the ocean

We expected that the capacity to store PHAs would be common in Arctic bacterial taxa as it is commonly reported that the ability to store PHAs is widespread across prokaryotes^8^. Our results show a large diversity of taxa with the ability to polymerize PHAs as evidenced by the large number of genes clusters involved in PHA biosynthesis. The phylogenetic restriction of PHA metabolism within *Alpha-* and *Gammaproteobacteria* that we observe in our data is consistent with what has been reported in other world oceans^68,69^. However, taxa from marine environments reported to store PHAs were restricted to those that could be cultivated^68^. To our knowledge, no studies attempted to systematically assess the capacity of ocean microbiomes to synthesize PHAs. Our work is the first report to systematically survey the capacity of ocean microbiomes to synthesize PHAs. Our study can therefore serve as a basis to further explore PHA storage capacities in the global ocean microbiome.

Our study considerably expands the known diversity of bacterial taxa potentially storing TAGs in the ocean. Most of the bacterial taxa with this ability were previously identified in C-rich environments such as municipal and agro-industrial waste waters^70,71^, or in environments with long periods of C scarcity such as desert soils^5,50^. These taxa are largely within the *Actinobacteria* and *Gammaproteobacteria* groups ^9,72,73^ but genes coding for the DGAT were also found in members of *Alpha-, Beta- and Deltaproteobacteria* as well as in *Bacteroidetes*^74^. In the ocean, TAG biosynthesis has frequently been reported for bacterial species of the *Gammaproteobacteria* genera *Alcanivorax*^75^ and *Marinobacter*^76^ in oil spills^77–79^. The diversity of taxa able to store TAGs that we report in this study was therefore unexpected. In addition to the expected *Actinobacteria*, we also report genomes in the group *Myxoccocota, Planctomycetota, Verrucomicrobia, Acidobacteria, Desulfobacterota, Chloroflexota and Latescibacteria*. A *Myxoccocota* species has been shown to produce TAGs^80^ and TAG biosynthesis was identified in a *Planctomycete*^81^. However, to our knowledge, our study considerably expands the diversity of marine bacterial taxa known to synthesize TAGs in the ocean. Further experimental work, including microscopy observations, will be needed to confirm the accumulation of TAGs in these groups within the Arctic Ocean.

### The ecology of NL storage for the Arctic Ocean heterotrophic prokaryotes

The storage of NLs may be used by a subset of the Arctic Ocean bacterial population as a life strategy to survive the highly variable seasonal conditions of the upper water column of the Arctic Ocean. The storage of compounds in summer and its mobilization in winter is a strategy that has been observed in soil bacteria to survive through winter^45^ but has not been reported in the Arctic Ocean so far. In addition, NLs may help bacterial cells resist other stressors. As highly reduced compounds, PHAs help maintain the redox state of the cell under oxidative stress^12,82^. PHAs could therefore participate to protect the Arctic Ocean bacteria from cold-induced oxidative stress, similarly to what has been shown in the Antarctic bacterium *Pseudomonas* sp. 14-3^83,84^. In the upper water column of the Arctic Ocean (surface, SCM, FDOMmax), bacteria probably accumulate NLs slowly under a *reserve storage* scenario. Under that scenario, the accumulation of reserves competes with other metabolic processes and would be slow. Evidence of NL *reserve storage* includes *Pseudomonas putida* accumulating PHA up to 26% of dry cell mass^85^ and *Rhodococcus opacus* accumulating TAG up to 21 % of its dry cell mass^86^ both in C-limited conditions. The low abundance of MAGs able to store TAGs or PHAs in the surface samples, where phytoplankton blooms occur, suggests that the storage of NLs in bacteria does not rely on C sources from phytoplankton blooms. Rather, in the upper water column, bacterial storers are most abundant in the SCM and the FDOMmax. A particularity of these water features is the accumulation of terrestrial organic matter whose concentration stays relatively stable throughout the year. This terrestrial organic matter may feed the *reserve storage* of NLs in bacteria. Supporting this hypothesis, we observed an enrichment of aromatic compound degradation genes in NL storage MAGs found in the SCM and FDOMmax, a metabolic process linked to the degradation of terrestrial organic matter in the Arctic Ocean^87^.

In the deep water of the Arctic Ocean and other oceans, bacteria may rather store NLs under a *surplus storage* scenario. MAGs with the capacity to synthesize TAGs that were abundant in deep samples possessed more CAZymes genes. This may reflect the capacity to recycle carbohydrates from dying bacteria or phytoplankton-derived marine snow^88^. The carbon and nutrient concentration of marine snow and dying bacteria are up to 4 orders of magnitude more elevated than the background water concentrations^89^. An encounter of a marine snow particle or a dying bacterium may therefore represent a *surplus storage* scenario for the taxa able to store TAGs. They may use the TAGs accumulated during these encounters to survive until the next encounter. Interestingly, studies showed that storage of TAGs is favoured over PHAs in cyclic feast/fast conditions when there is an overlap in time between the supply of C and N during the feast stage^90^. As marine snow and dying bacteria also contain nitrogen, this may explain why we preferentially find TAGs and not PHAs biosynthesis genes in MAGs that are abundant in deep samples.

Throughout the water column, regardless of the storage scenario, the storage of NLs may be favored by strategies limiting the energy demand of NL biosynthesis. Using exogenous FAs instead of synthesizing FAs to incorporate FAs into NLs greatly reduces the energy cost of NL storage. Our results show that a higher percentage of MAGs able to store TAGs and PHAs contained genes involved in exogenous FA import and use compared to MAGs with no NL metabolism genes. Exogenous FAs can be incorporated directly into TAGs. FAs can also be incorporated into PHAs either by being diverted from β-oxidation (to form medium-chain length PHA) or by going through β-oxidation and producing acetyl-CoA that can be used to synthesize polyhydroxybutanoate (short-chain PHA). The absence of the medium-chain length PHA hydrolase gene in our data suggests that the second option is preferred.

### A possible role for prokaryotic NL storage to sustain the winter Arctic Ocean ecosystem

We propose the hypothesis that NL storage in the upper water column of the Arctic Ocean shifts from phototrophic eukaryotic based production from spring through fall towards heterotrophic prokaryotic based production during the winter polar night (**Figure 7**). There is evidence that the Arctic Ocean is not an unproductive desert during the polar night but still retains significant levels of biological activity^3,91^. It implies some light independent levels of metabolic activity at the base of the food web, hence in the microbiomes^92^. The storage of PHAs and TAGs by the bacterial taxa during the spring, summer, fall and throughout winter may participate significantly to the carbon pool necessary to maintain this level of metabolic activity through the polar night. The C stored in PHAs and TAG-accumulating bacteria could then go up and support the food chain through bacterivory. Bacterivory is a common strategy for phototrophic eukaryotes to overwinter in polar seas^93^. Reports have shown that, in the winter, phototrophic *Micromonas pusilla*, one of the most abundant phototrophs in the Arctic Ocean, can survive by grazing on bacteria^94,95^ and that winter prokaryotic populations are controlled by grazing^96^. This suggests a possible new and important role for the NL storage metabolism in the Arctic prokaryotic communities in sustaining the ecosystem during the polar night.

**Figure 7:**
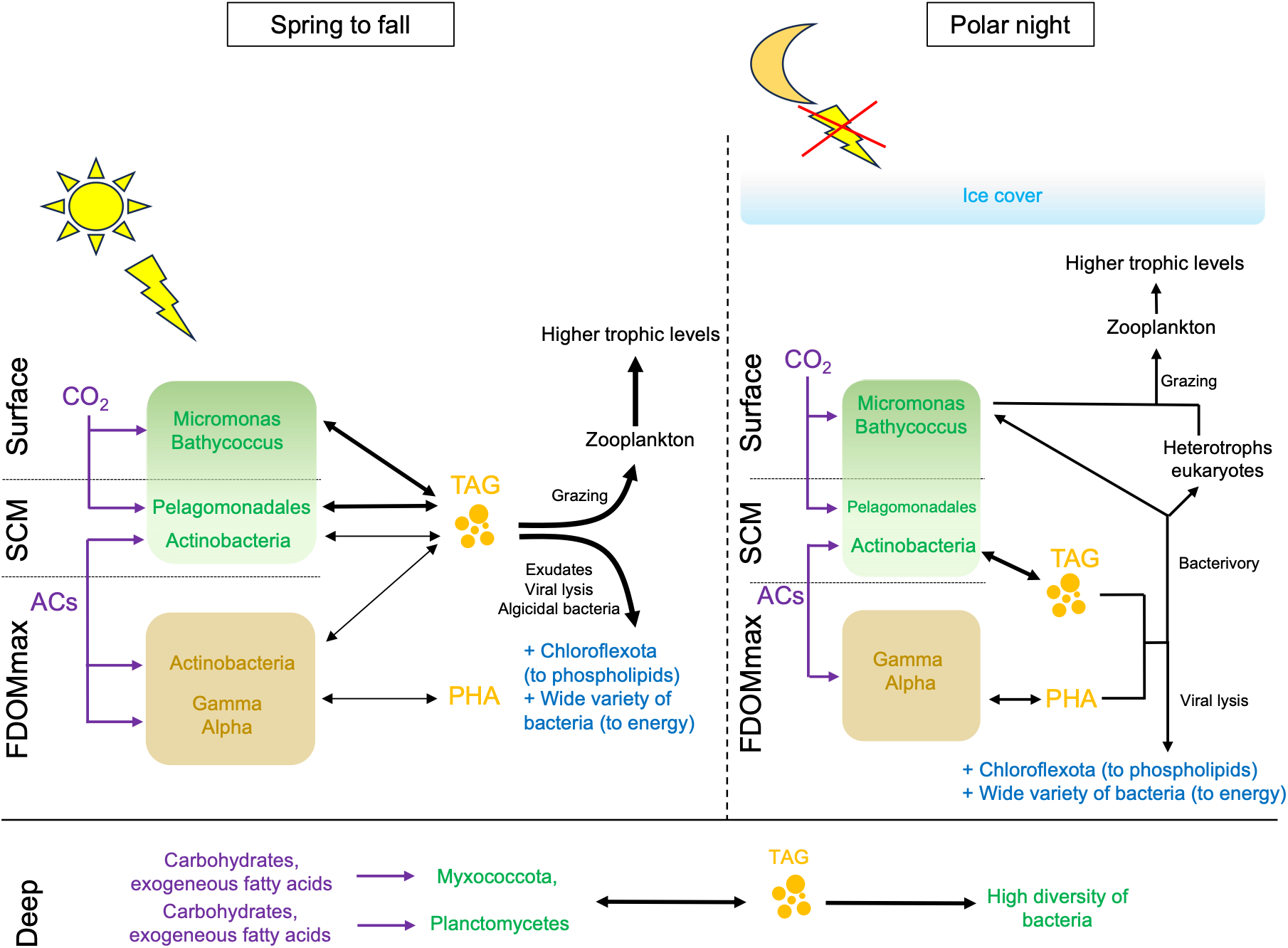
Proposed model for the metabolism of neutral lipids in the microbiomes of the Arctic Ocean.

## Conclusion

The Arctic Ocean is characterized by an intense seasonality that drives strong changes in conditions, nutrients, and energy availability for the microbiomes. In this context, the accumulation of NLs as energy and carbon reserves may play an important role as a strategy to survive through periods of resource scarcity during winter. Given that lipids from microbial origin are an important growth resource for the Arctic Ocean food chain, and that NLs may constitute a significant fraction of microbial lipids, it is crucial to understand the metabolism of neutral lipids in the Arctic Ocean microbiome. In this study, we showed that the prokaryotic and picoeukaryotic communities of the Arctic Ocean photic were enriched in TAG biosynthesis genes due to the dominance of eukaryotes in the phytoplankton fraction. We showed that these TAGs can be transferred and consumed by a “lipotrophic” fraction of the heterotrophic prokaryotes, shedding light on a new important process of the C cycle in the Arctic Ocean. We also discovered bacterial taxa able to synthesize NLs by using aromatic compounds from the disproportionately high fraction of terrestrial organic matter that washes off in the Arctic Ocean. Altogether, these results unravel the importance of NL metabolism for the adaptation of the microbiomes in the Arctic Ocean, but also postulates a new significance for microbial NL metabolism for the Arctic Ocean C cycle and in supporting the marine food web.

## Acknowledgments

The data were collected aboard the CCGS Louis S. St-Laurent in collaboration with researchers from Fisheries and Oceans Canada at the Institute of Ocean Sciences and Woods Hole Oceanographic Institution’s Beaufort Gyre Exploration Program and are available at http://www.whoi.edu/beaufortgyre. We would like to thank both the captain and crew of the CCGS Louis S. St-Laurent and the scientific teams aboard. We also thank Québec-Océan for their scientific and financial support.

## Declarations

### Contributions

TG and DAW designed the study. TG generated the metagenomic data and performed the bioinformatic analyses. TG wrote the manuscript and DAW commented and edited the manuscript. SM performed the binning and assembly of MAGs. VEO performed the functional annotation of the MAGs. All the authors read and approved the final manuscript.

### Ethics approval and consent to participate

Not applicable

### Competing interests

The authors declare that they have no competing interests.

### Availability of data and materials

The metagenomic data generated in this study are available in the Integrated Microbial Genomes database at the Joint Genome Institute at https://img.jgi.doe.gov/cgi-bin/m/main.cgi?section=GenomeSearchList&page=displayTaxonList&searchFilter=all&searchTerm=Gs0134626&file=all761564&allDataFiltersFile=allGenomeDataFilters761564, GOLD Project ID: Gs0134626. Metagenome-assembled genome projects will be deposited at Dryad (https://datadryad.org/)

## Notes

### Competing Interest Statement

The authors have declared no competing interest.

### Summary of Updates

Layout of the manuscript file. Notes were still present on the manuscript, we removed the notes.

