## Supplementary figures and table legends for "Metagenomic Discovery of Neutral Lipid Metabolism Pathways in the Arctic Ocean Microbiomes Suggests a Potential New Role in Survival and Oceanic Carbon Cycling"

***Authors and affiliations***

1 - Department of Biology, Concordia University, 7141 Sherbrooke St. West, Montreal, QC, H4B 1R6, Canada

2 - Division of Science, New York University Abu Dhabi, Saadiyat Marina District, Abu Dhabi, United Arab Emirates

### Figures

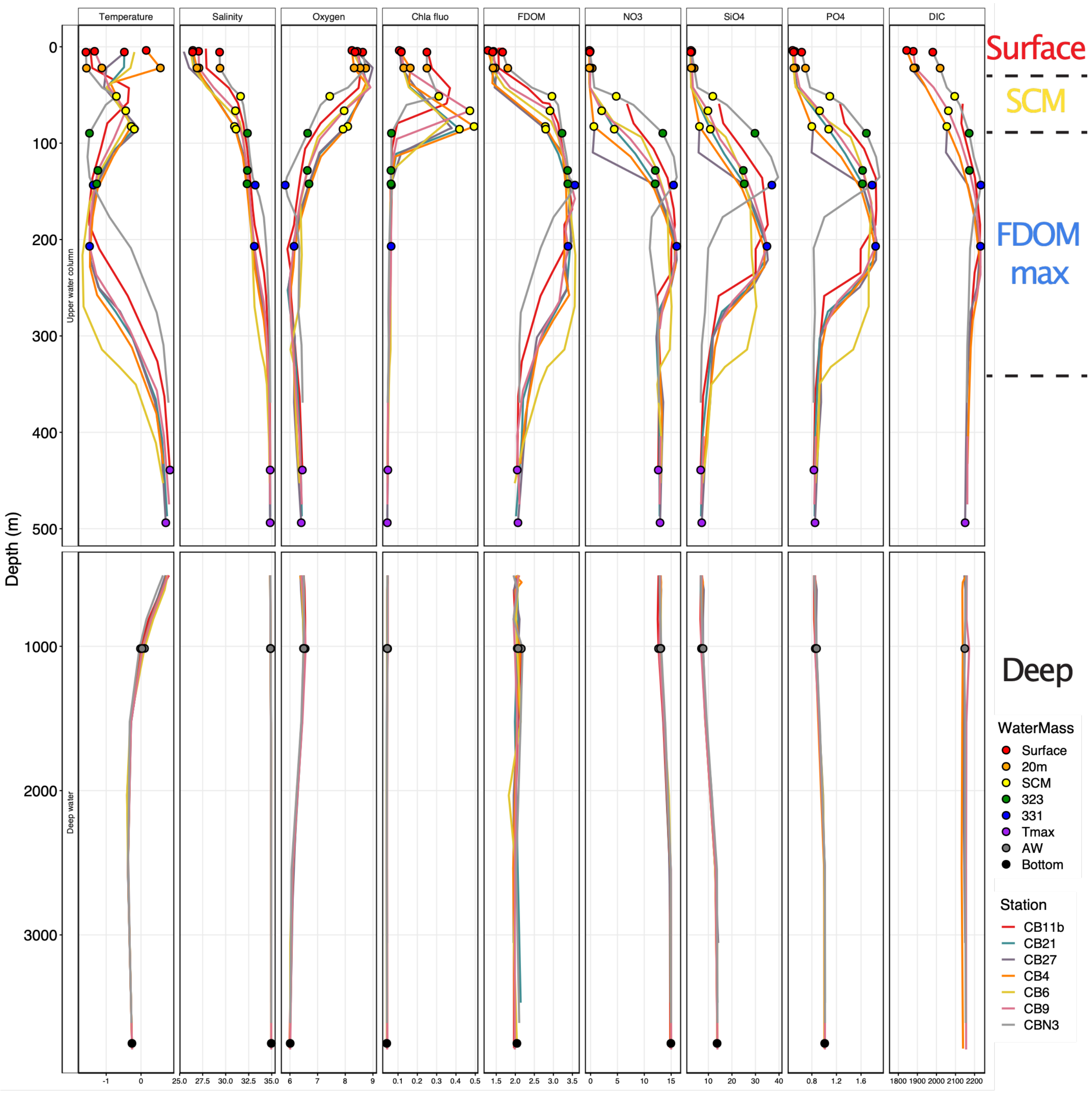


**Figure A1:** Depth profiles of environmental variables for the 7 stations sampled in the Canada Basin for this study. Points on the profiles represent the location of the metagenomes sampled.

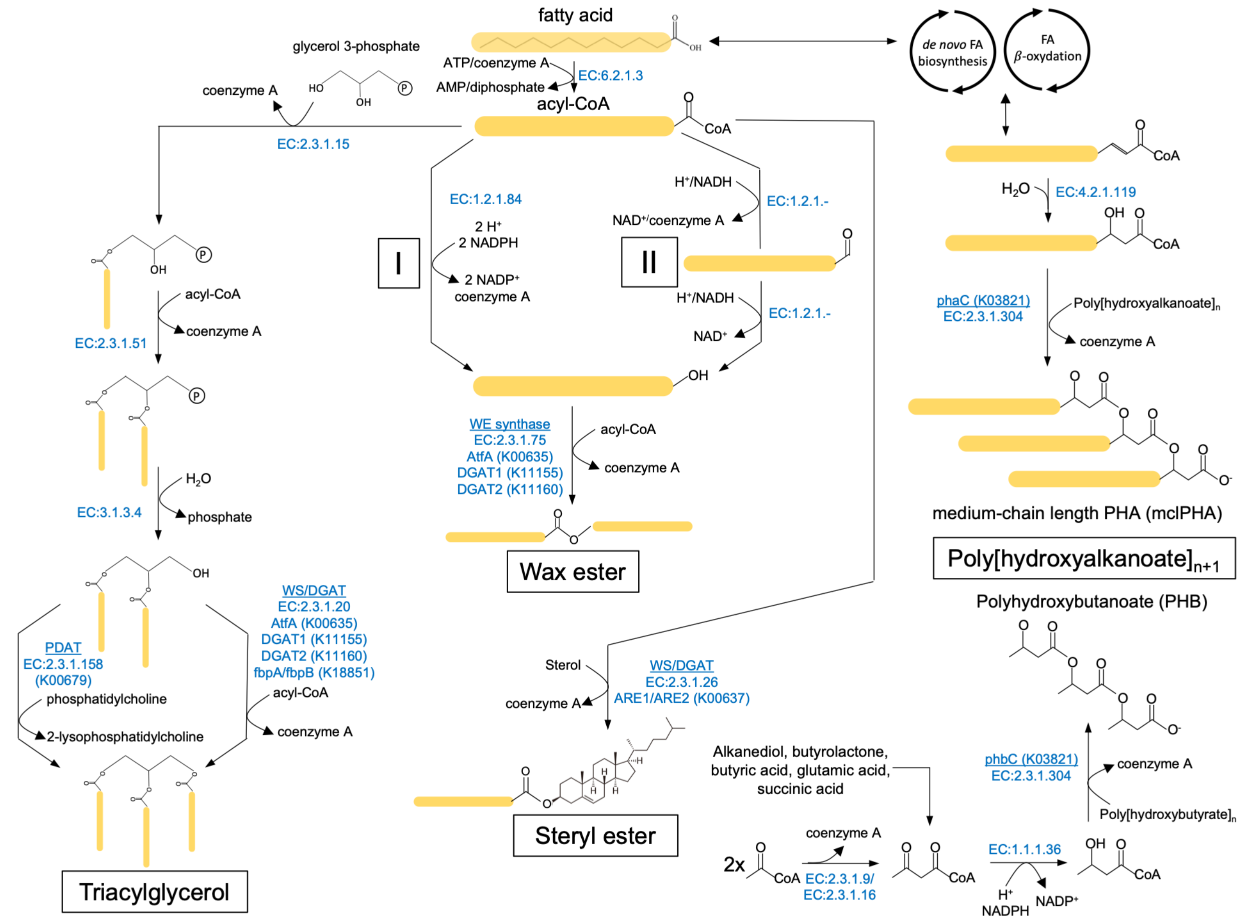


**Figure A2:** Detailed metabolic pathways for the biosynthesis of neutral lipids

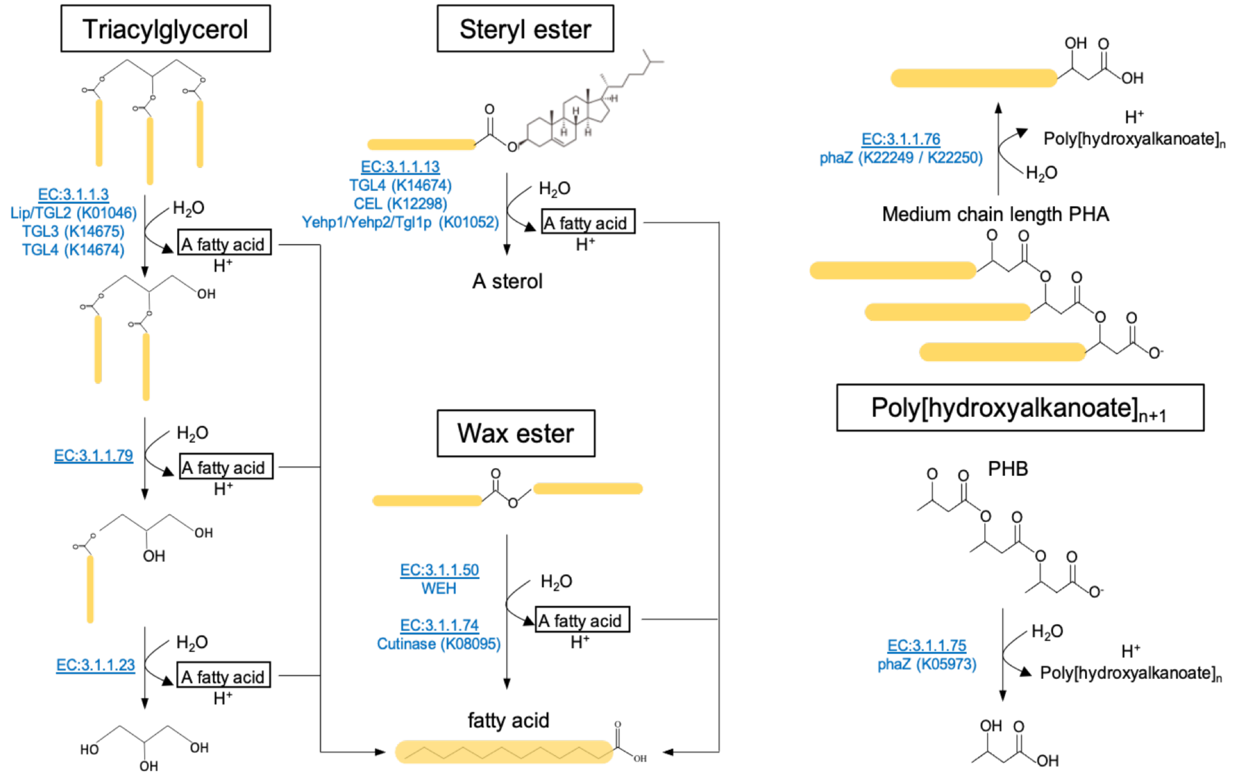


**Figure A3:** Detailed metabolic pathways for the degradation of neutral lipids

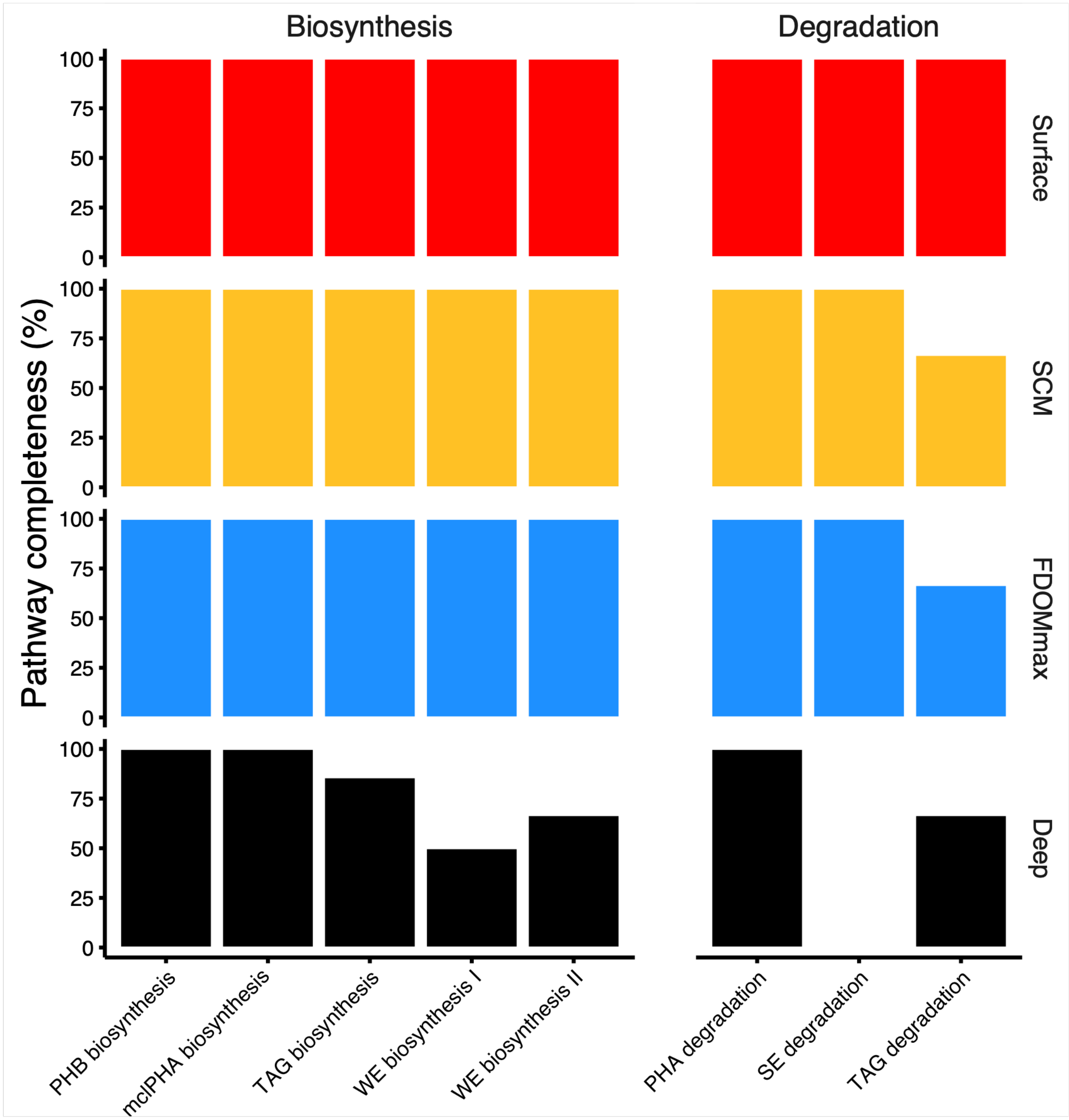


**Figure A4:** Completeness of the neutral lipid biosynthesis and degradation pathways within the microbiome metagenomes of the different water column features of the Canada basin

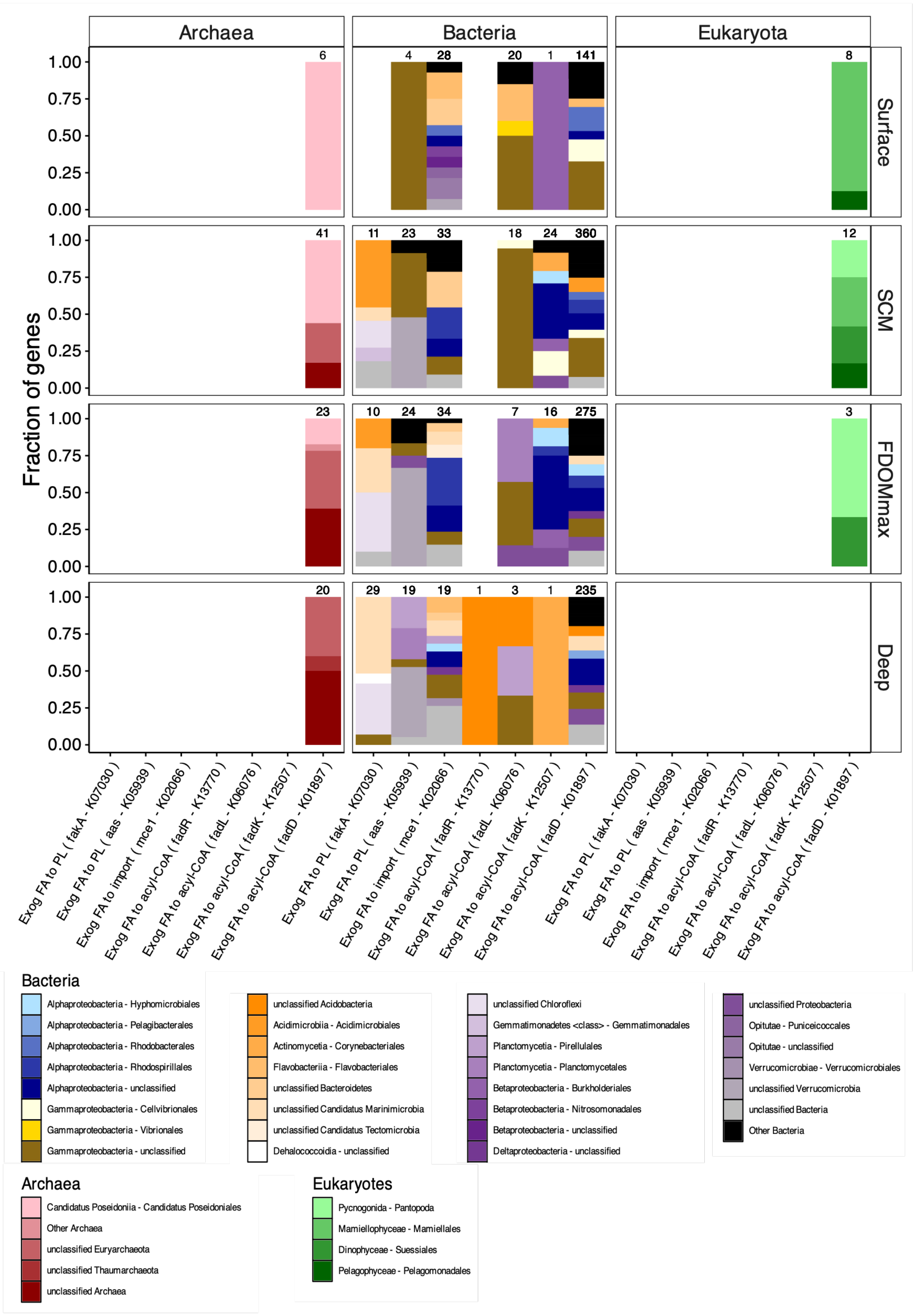


**Figure A5:** Taxonomic identity of genes involved in the import of exogeneous fatty acids for the microbiomes of the Canada basin

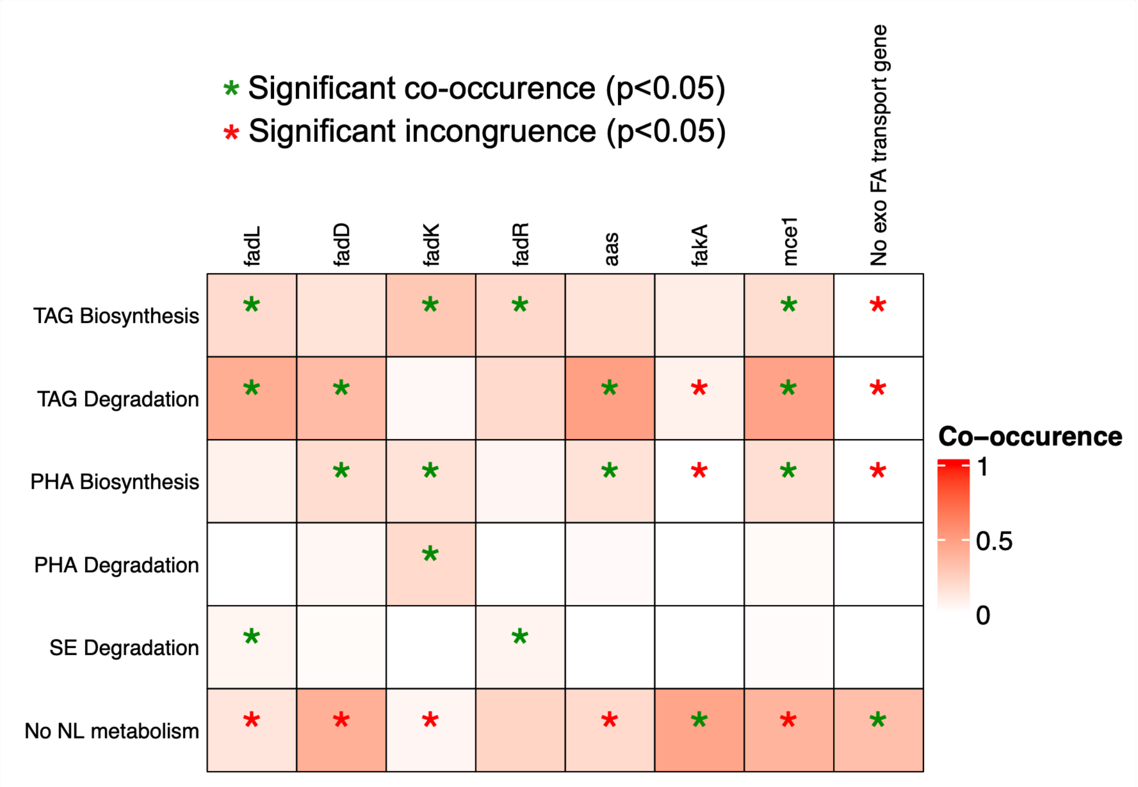


**Figure A6:** Co-occurrence of genes involved in NL metabolism and exogenous FA import in our MAG dataset. The co-occurrence is calculated based on the Sorensen index. Green stars indicate statistically significant co-occurrence of genes in MAGs. Red stars indicate a significant absence of one gene if the second gene is present (incongruence of genes).

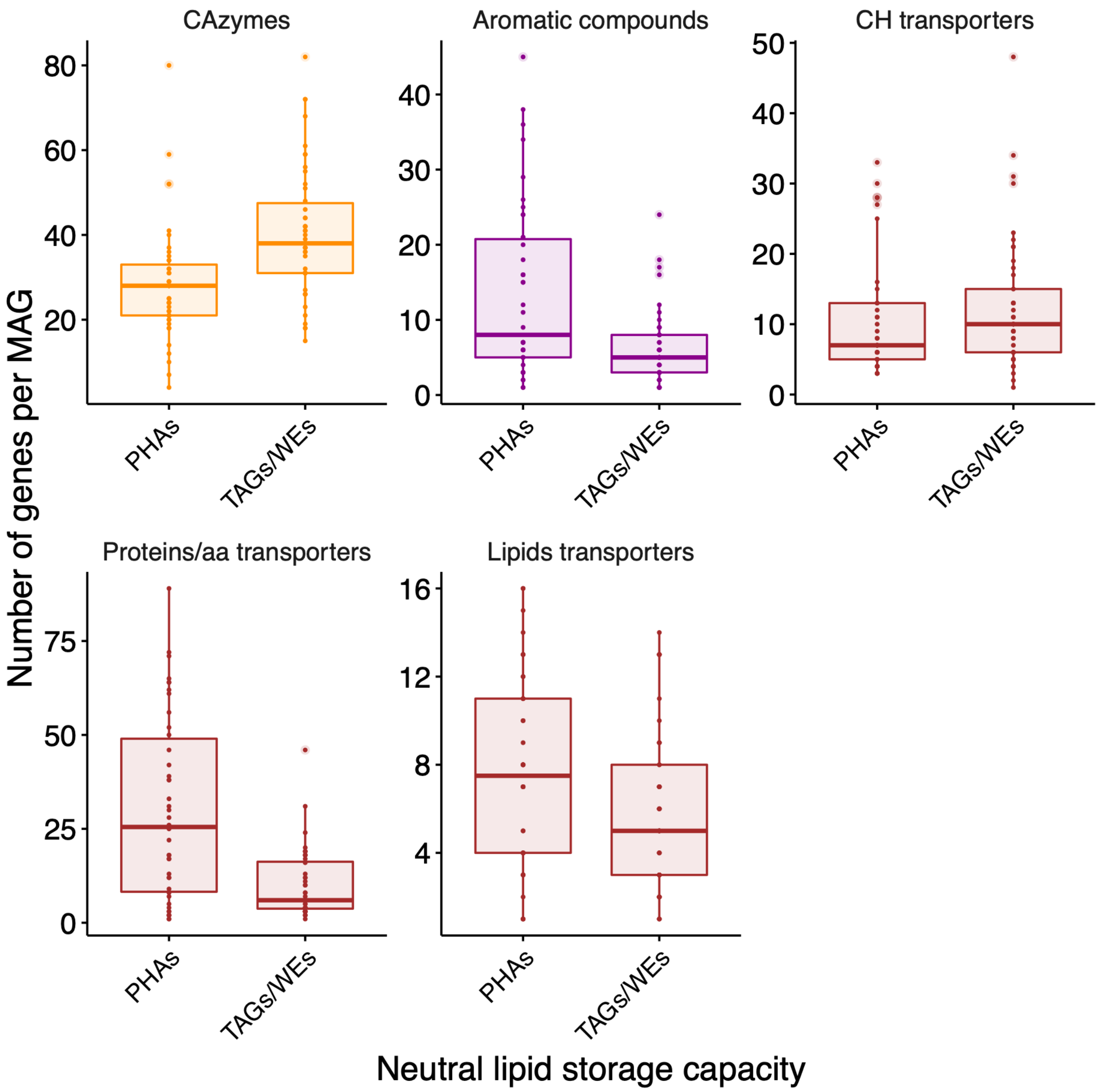


**Figure A7:** Comparison of the number of genes associated with various metabolic processes (aromatic compound degradation, sugar utilization, protein and lipid transporters) between MAGs with TAG and PHA biosynthetic capacity.

#### Appendix tables

**Table A1:** Data availability, metadata and physicochemical properties of the samples collected for this study.

**Table A2:** List of metabolic pathways (obtained from metacyc, https://metacyc.org/) involved in the biosynthesis and degradation of neutral lipids as well as import of exogenous fatty acids. The corresponding pathway names used this study are specified. KO numbers, and gene name information is included for reactions identified as markers for these pathways.

**Table A3:** Information and sources of publicly available metagenomes from other studies used in this study.

**Table A4:** List of CAZymes used in the analysis of this study and their corresponding EC numbers.

**Table A5:** List of aromatic compound degradation pathways and the EC numbers that belong to each pathway.

**Table A6:** List of all the transporter genes retrieved from the Kyoto Encyclopedia of genes and genomes (https://www.genome.jp/kegg/), as well as their KEGG ortholog number, their substrate type, and their pathway hierarchy.

**Table A7:** Assemblies of metagenomes used to generate the metagenome-assembled genomes of this study.
